# MmpL3 is the flippase for mycolic acids in mycobacteria

**DOI:** 10.1101/099440

**Authors:** Zhujun Xu, Vladimir A Meshcheryakov, Giovanna Poce, Shu-Sin Chng

## Abstract

The defining feature of the mycobacterial outer membrane (OM) is the presence of mycolic acids (MAs), which in part render the bilayer extremely hydrophobic and impermeable to external insults, including many antibiotics. While the biosynthetic pathway of MAs is well studied, the mechanism(s) by which these lipids are transported across the cell envelope is(are) much less known. MmpL3, an essential inner membrane (IM) protein, is implicated in MA transport, but its exact function has not been elucidated. It is believed to be the cellular target of several anti-mycobacterial compounds; however, evidence for direct inhibition of MmpL3 activity is also lacking. Here, we establish that MmpL3 is the MA flippase at the IM of mycobacteria, and is the molecular target of BM212, a 1,5-diarylpyrrole compound. We develop assays that selectively access mycolates on the surface of *Mycobacterium smegmatis* spheroplasts, allowing us to monitor flipping of MAs across the IM. Using these assays, we establish the mechanism-of-action of BM212 as a potent MmpL3 inhibitor, and employ it as a molecular probe to demonstrate the requirement for functional MmpL3 in the transport of MAs across the IM. Finally, we show that BM212 binds MmpL3 directly and inhibits its activity. Our work provides fundamental insights into OM biogenesis and MA transport in mycobacteria. Furthermore, our assays serve as an important platform for accelerating the validation of small molecules that target MmpL3, and their development as future anti-tuberculosis drugs.

## Significance statement

Biological membranes define cellular boundaries, allow compartmentalization, and represent a prerequisite for life; yet, our understanding of membrane biogenesis remains rudimentary. Mycobacteria, including the human pathogen *Mycobacterium tuberculosis*, are surrounded by a double-membrane cell envelope that makes them intrinsically resistant to many antibiotics. Specifically, the outer membrane contains unique lipids called mycolic acids, whose transport mechanism across the envelope is unknown. In this study, we established the role of an essential membrane protein as the flippase for mycolic acids, and demonstrated that this protein is a direct target of an anti-mycobacterial compound. Our work provides insights into outer membrane biogenesis and lipid transport in mycobacteria, and the means to evaluate drugs that disrupt mycolic acid transport at the inner membrane.

The outer membrane (OM) of *Mycobacterium tuberculosis*, the causative agent of tuberculosis (TB), is distinctively characterized by the abundance of mycolic acids (MAs), C_60_-C_90_ long chain, branched fatty acids, packed together to produce a bilayer with markedly reduced fluidity and permeability (1). These MAs come in the forms of trehalose monomycolates (TMMs), trehalose dimycolates (TDMs), and mycolates covalently attached to arabinogalactan (AG) polysaccharides, which are in turn linked to the peptidoglycan and collectively known as the mAGP complex (Fig. 1*A*). MAs are synthesized at the inner membrane (IM) as TMMs via a highly-conserved and well-characterized pathway (2), which is the target of first line anti-TB drug, isoniazid (3). How MAs are transported across the cell envelope and assembled into the OM, however, is less understood; proteins mediating TMM flipping across the IM and transit across the periplasm have not been identified and/or characterized (Fig. 1*A*). At the OM, the Ag85 complex transfers a mycolate chain from one TMM molecule to another to form TDM, or to the AG polysaccharides to form the mAGP complex (4). Tethering the OM to the cell wall via the AG polysaccharides further rigidifies the membrane, making it extremely impermeable to a wide range of compounds, including many antibiotics (1). The OM, and hence MAs, are essential for mycobacterial growth.

**Fig. 1.**
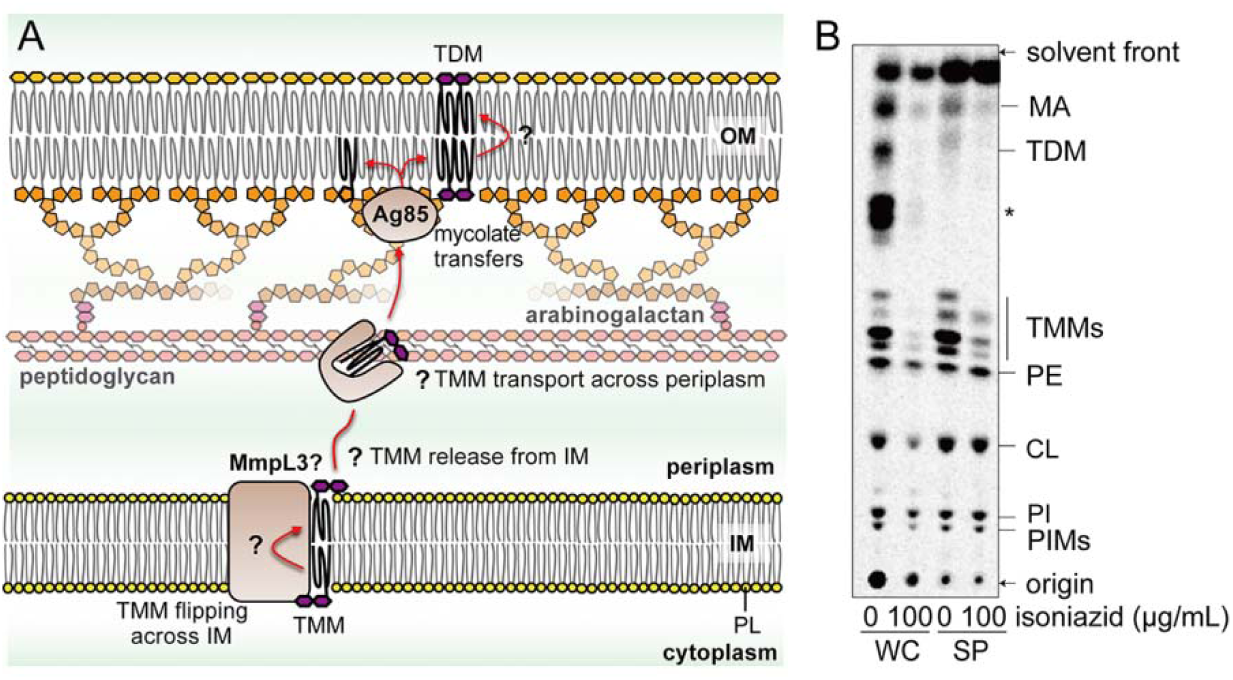
TMM biosynthesis is intact in mycobacterial spheroplasts. (*A*) A schematic diagram illustrating the processes important for MA transport across the cell envelope. Following synthesis, TMMs must be flipped across the IM, released from the IM and then transported across the periplasm (presumably via a chaperone). MmpL3 is implicated in TMM transport at the IM, but its exact role has not been elucidated. At the OM, the Ag85 complex transfers the mycolate chain from TMM to cell wall-linked AG polysaccharides, or to another TMM to form TDM. Other known lipid species found in the OM and IM are omitted for simplicity. PL, phospholipid. (*B*) TLC analysis of newly-synthesized [^14^C]-labelled lipids extracted from wild-type *M. smegmatis* cells (WC) and spheroplasts (SP), visualized by phosphor imaging. Lipids were radiolabelled in the presence or absence of isoniazid as indicated. The developing solvent system comprises chloroform-methanol-water (30:8:1). A mycolate-based species that appears only in the presence of glucose is indicated with an asterisk (*). PE, phosphatidylethanolamine; CL, cardiolipin; PI, phosphatidylinositol; PIM, phosphatidylinositol mannoside.

Recently, a conserved essential IM protein, MmpL3 (<u>M</u>ycobacterial <u>m</u>embrane protein Large 3), has been implicated in MA transport. Depletion of *mmpL3* in *Mycobacterium smegmatis* results in accumulation of TMMs and reduced formation of TDMs and AG-linked mycolates (5, 6), suggesting an impairment in TMM transport to the OM. Consistent with this, MmpL3, like other MmpL proteins, belongs to the resistance, nodulation, and cell division (RND) protein superfamily, and is believed to be a proton motive force (pmf)-dependent transporter (7). Based on its cellular localization, MmpL3 is likely involved in either TMM flipping across the IM, TMM release from the IM into the periplasm, or both (Fig. 1*A*). Yet, its exact role has not been clearly defined, due largely to the lack of functional assays for its putative transport activity. Treatment of mycobacteria with a few structurally-distinct small molecule scaffolds, including ethylenediamines (e.g. SQ109) (8), 1,5-diarylpyrroles (e.g. BM212) (9,10), adamantyl ureas (e.g. AU1235) (5), and others (11-15), result in similar changes in mycolate species as in *mmpL3* depletion. These compounds inhibit growth, and select for resistance mutations in *mmpL3*; however, there is limited evidence that they are direct MmpL3 inhibitors. The lack of activity assays for MmpL3 made it impossible to test the proposed mechanism-of-action of these putative inhibitors.

Here, we report that MmpL3 is the TMM flippase at the IM. Using a spheroplast model, we developed assays to monitor IM topology of TMM. We found that 1,5-diarylpyrrole BM212 inhibits TMM flipping across the IM in wild-type spheroplasts. Furthermore, we established that specific MmpL3 variants confer resistance against this inhibition, indicating that MmpL3 is required for flipping TMM across the IM. Finally, we demonstrated that BM212 binds MmpL3 in vitro in a specific manner, and therefore directly targets MmpL3. Our work represents the first demonstration of lipid transport activity of a key member of the MmpL protein family, and highlights the importance of using small molecule probes to interrogate protein function. Our assays have great utility in the validation and development of MmpL3-targeting small molecules as future anti-TB drugs.

## Results

### Spheroplasts serve as a viable system to monitor TMM topology

To develop a functional assay for TMM flipping across the IM, we sought a system where TMM topology in the IM can be monitored. Mycobacterial spheroplasts are ideal for this purpose because they lack the OM and cell wall (16), and are bound only by the IM (17, 18), providing easy access to molecules-of-interest at this membrane. Due to the loss of periplasmic contents upon the formation of spheroplasts, we also expect the transport pathway(s) for TMM to the OM to be disrupted, thereby resulting in accumulation of TMM at the IM. *M. smegmatis* spheroplasts were successfully generated via sequential treatment with glycine and lysozyme (SI Appendix, Fig. S1), as previously reported (16). To examine whether MA synthesis is intact in spheroplasts, we profiled newly-synthesized lipids metabolically labelled with [^14^C]-acetate. Thin layer chromatographic (TLC) analysis of lipids extracted from whole cells revealed a few major species whose syntheses are inhibited by isoniazid, indicating that these are mycolate-based lipids (Fig. 1*B*). We assigned two of these species as TDM and TMM on the basis of reported retention factors of these lipids on TLC plates developed under the same solvent system (19). We showed that mycolates are still produced in *M. smegmatis* spheroplasts; however, the extracted lipids only contain TMM, but not TDM. Furthermore, we can no longer detect newly-synthesized mAGP in the form of liberated mycolic acid methyl esters (MAMEs) in these spheroplasts (SI Appendix, Fig. S2). These results are consistent with the loss of Ag85 enzymes and the OM, where TDM and mAGP syntheses occur, and also with the lack of TMM transport to any possible remnants of the OM. Given the extreme hydrophobicity of mycolates, we conclude that newly-synthesized TMMs accumulate in the IM of spheroplasts, thus establishing a platform for monitoring TMM flipping across the bilayer.

### TMMs accumulated in spheroplasts reside in the outer leaflet of the IM

We next examined whether newly-synthesized TMMs accumulated in the inner or outer leaflet of the IM in spheroplasts, by monitoring its accessibility to degradation by recombinant LysB, a lipolytic enzyme. LysB is a mycobacteriophage-encoded esterase that is specific for mycolates and plays the role of an endolysin important for the release of phage particles from infected cells (20, 21). Substantial amounts (˜77%) of newly-synthesized TMMs in spheroplasts are readily and specifically hydrolyzed by purified LysB with the concomitant release of MAs (Fig. 2*A*). Part of this hydrolysis can be attributed to the background exposure of TMMs to LysB in a subset of spheroplasts that lysed during the experiment (˜30-50% cell lysis irrespective of the addition of LysB; Fig. 2*B*, and SI Appendix, Fig. S3). The rest of newly-synthesized TMMs that were cut by LysB are likely accessible on the surfaces of intact spheroplasts. We showed that an inactive LysB variant does not result in the same effect (Fig. 2*A*). In addition, we demonstrated that LysB does not enter intact spheroplasts (Fig. 2*B*), nor does it induce additional cell lysis compared to controls (Fig. 2*B*, and SI Appendix, Fig. S3). Taken together, these results establish that most of newly-synthesized TMMs have been translocated across the IM in intact spheroplasts, and therefore reside in the outer leaflet of the membrane.

**Fig. 2.**
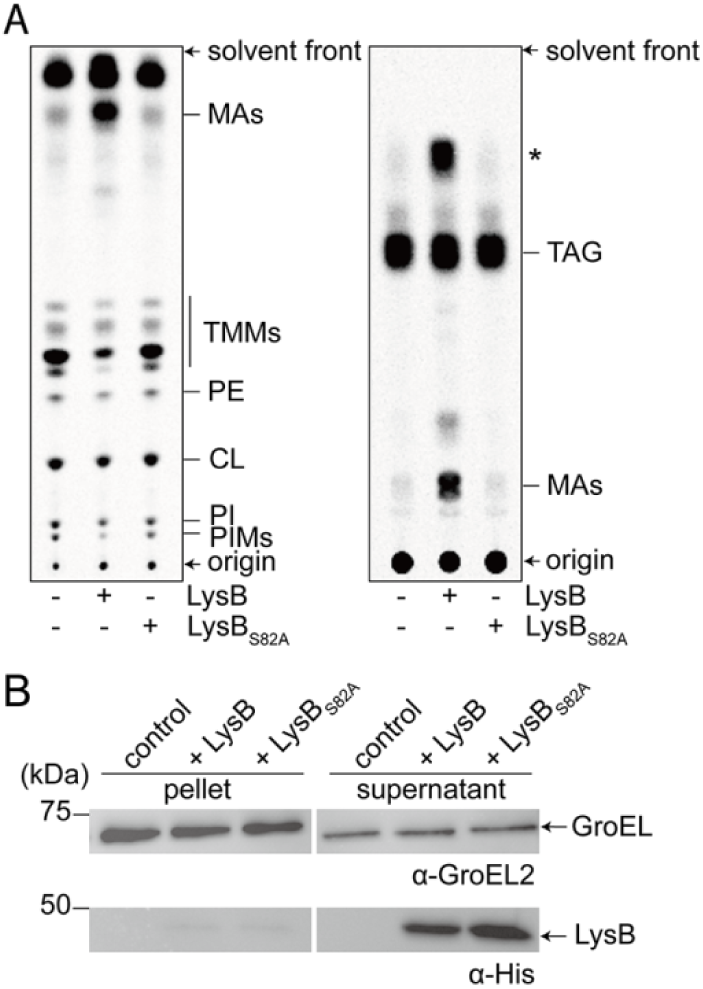
Newly-synthesized TMMs in mycobacterial spheroplasts are accessible to degradation by LysB, indicating that these TMMs reside in the outer leaflet of the IM. (*A*) TLC analyses of newly-synthesized [^14^C]-labelled lipids extracted from *M. smegmatis* spheroplasts treated with functional or non-functional (S82A) LysB. Lipids were resolved on TLCs developed using solvent systems comprising either chloroform-methanol-water (30:8:1) (*left*) or hexane-diethylether-acetic acid (70:30:1) (*right*), followed by phosphor imaging. In addition to MA, treatment with functional LysB also resulted in the release of an unidentified apolar lipid, annotated with an asterisk (*). TAG, triacylglycerol. (*B*) a-GroEL2 and a-His immunoblot analyses of pellet and supernatant fractions obtained from sedimentation of *M. smegmatis* spheroplasts treated with functional or non-functional (S82A) LysB.

### MmpL3 is responsible for flipping TMM across the IM

Several compounds, including SQ109, BM212 and AU1235, are believed to affect MmpL3-mediated TMM transport because mutations in *mmpL3* confer resistance against these small molecules (5, 8, 9). While it is not yet clear if these compounds directly inhibit MmpL3, they may be useful as probes to determine if MmpL3 is responsible for TMM flipping across the IM. Specifically, we asked whether these small molecules are able to inhibit TMM flipping in wild-type spheroplasts, and whether they would become less effective in doing so in spheroplasts expressing MmpL3 variants that confer resistance against them. We first tested the effects of these compounds in our LysB assay in wild-type spheroplasts. Remarkably, BM212 and AU1235 are able to reduce LysB-mediated hydrolysis of newly-synthesized TMMs in *M. smegmatis* spheroplasts (Fig. 3*A* and *C*), at concentrations 2x and 4x above their reported minimal inhibitory concentrations (MICs) (SI Appendix, Table S1) (5, 9). In contrast, SQ109 has no effect (Fig. 3*B* and *C*). For BM212, hydrolysis by LysB is strongly reduced, close to the level of background attributed to non-specific spheroplast lysis (SI Appendix, Fig. S3). We showed that the effects of BM212 and AU1235 are not due to direct inhibition of LysB activity, since TMMs are still hydrolyzed in detergent-solubilized samples (SI Appendix, Fig. S4). Instead, significant amounts of newly-synthesized TMMs are no longer accessible to LysB in the presence of either compound in intact spheroplasts, indicating inhibition of TMM flipping across the IM. As an alternative method to assess TMM topology in the IM and the effects of these drugs, we also examined the ability of membrane-impermeable fluorophore-conjugated streptavidin to bind to newly-synthesized TMMs engineered to contain biotin (biotin-TMMs) (Fig. 4*A*). Here, we metabolically labelled TMMs with 6-azido-trehalose (22), which allowed us to covalently attach an alkyne-containing biotin probe to TMM via the bio-orthogonal click reaction (23). In wild-type spheroplasts, biotin-TMMs can be detected on the surface, indicating that 6-azido-TMMs have been translocated across the IM (Fig. 4*C* and *F*). Biotin labelling of 6-azido-trehalose groups, and therefore detection with streptavidin, is largely prevented if LysB was added (SI Appendix, Fig. S5), confirming that we are visualizing trehalose-linked mycolates (i.e. TMMs) at the IM in spheroplasts. We demonstrated that both BM212 and AU1235 drastically reduce the amounts of 6-azido-TMMs, and hence biotin-TMMs, that can be labelled with streptavidin (Fig. 4*D, E* and *F*). Again, SQ109 is not effective in this assay (SI Appendix, Fig. S6). Consistent with the LysB accessibility assay, these results establish that BM212 and AU1235, but not SQ109, inhibit TMM flipping across the IM.

**Fig. 3.**
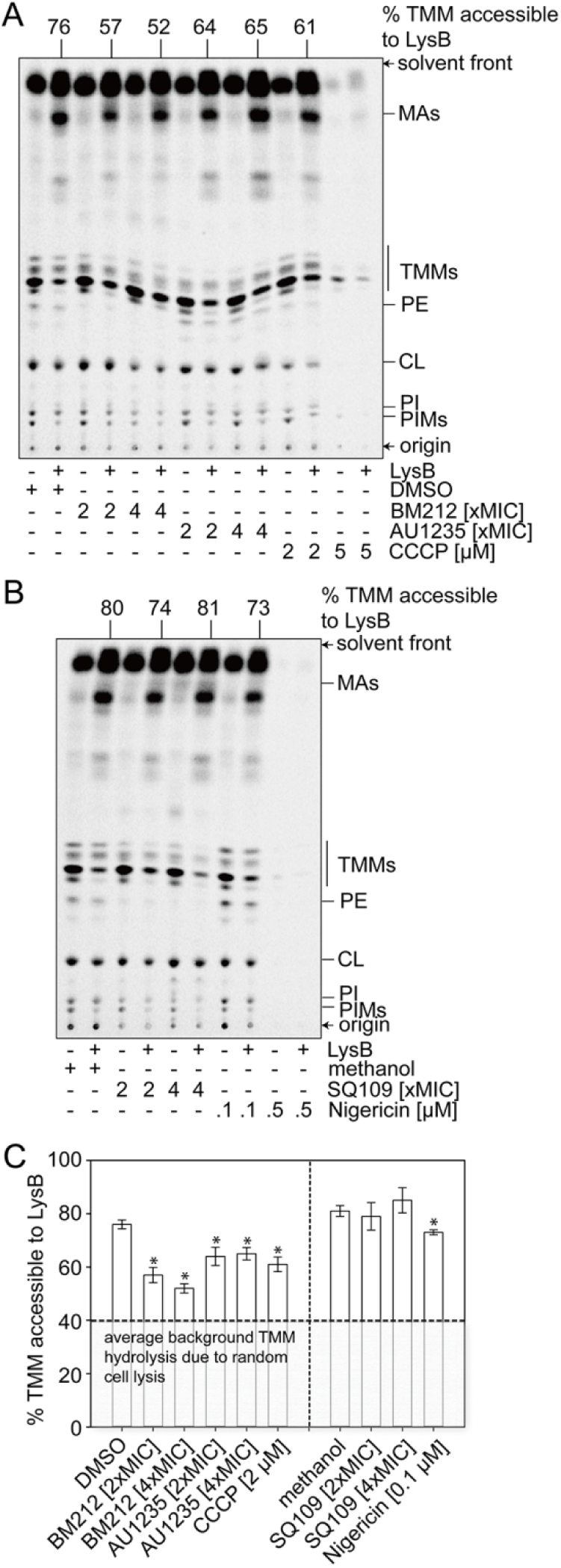
Anti-mycobacterial compounds, BM212 and AU1235, reduce TMM accessibility to LysB in spheroplasts, indicating inhibition of TMM flipping across the IM. (*A, B*) Representative TLC analyses of [^14^C]-labelled lipids newly-synthesized in the presence of indicated concentrations of (*A*) BM212, AU1235, and (*B*) SQ109, and extracted from *M. smegmatis* spheroplasts following treatment with or without purified LysB. The effects of pmf disruptors, carbonyl cyanide *m*-chlorophenyl hydrazone (CCCP) and nigericin, were also tested. At higher concentrations, these uncouplers affected lipid synthesis, consistent with the depletion of ATP. DMSO and methanol were used to dissolve the respective compounds and thus serve as negative controls. Equal amounts of radioactivity were spotted for each sample. The developing solvent system comprises chloroform-methanol-water (30:8:1). (*C*) A graphical plot showing the effects of various compounds on the amounts of LysB-accessible TMMs in spheroplasts. The percentage of TMMs accessible to LysB is given by the difference in TMM levels between samples with or without LysB treatment, normalized against that in control samples without LysB treatment. TMM levels in each sample were quantified as a fraction of total mycolates (TMM+MA). Average percentages and standard deviations from three biological replicates are plotted. The average background of TMM hydrolysis due to random cell lysis during the experiment (˜40%) is indicated. Student’s t-test: *, p < 0.05 compared to the corresponding DMSO or methanol controls.

**Fig. 4.**
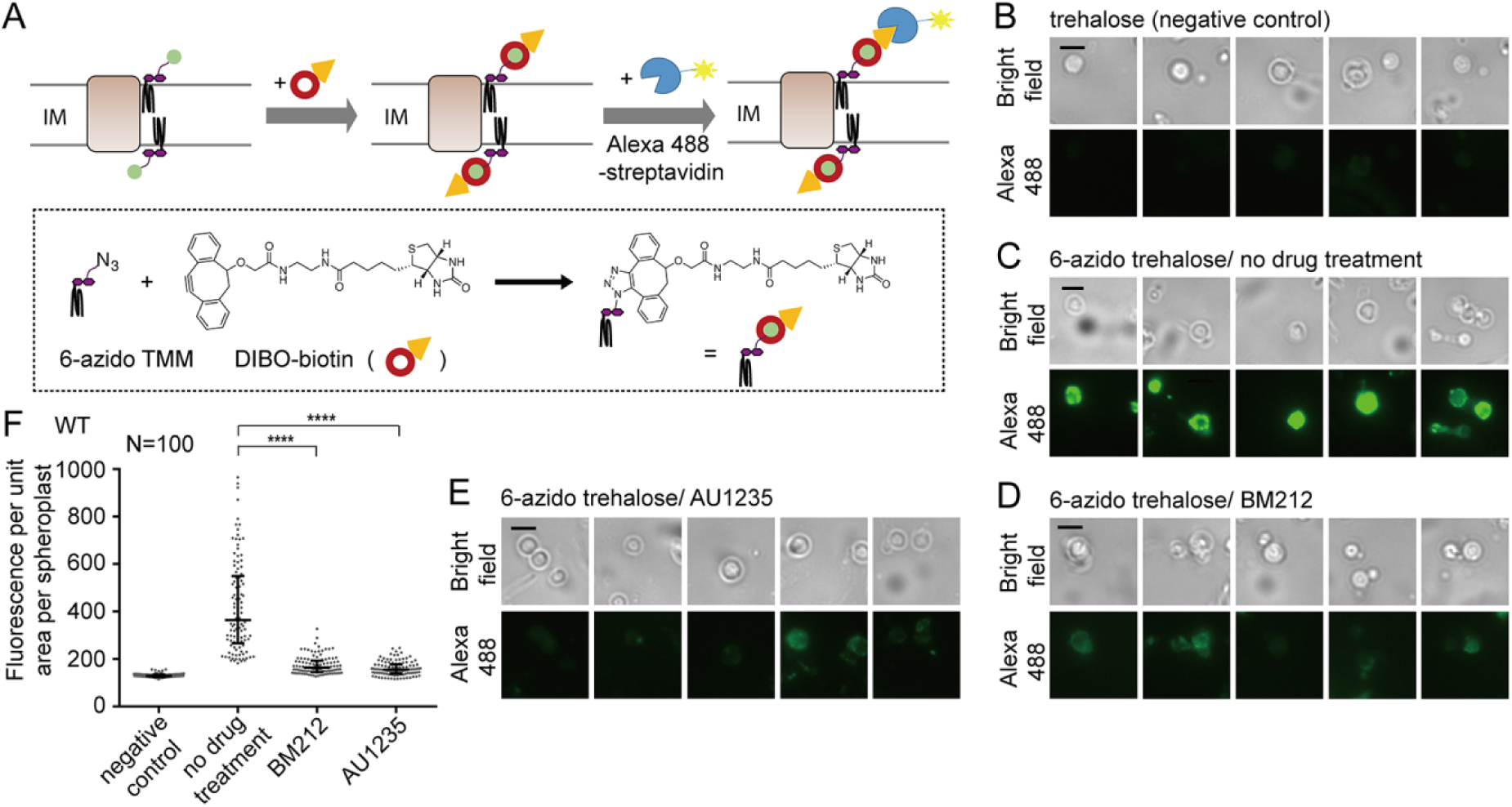
Anti-mycobacterial compounds, BM212 and AU1235, reduce surface display of 6-azido-TMMs in spheroplasts, indicating inhibition of TMM flipping across the IM. (*A*) A schematic diagram illustrating the 6-azido-TMM surface display assay. Spheroplasts were incubated with 6-azido-trehalose to allow synthesis of 6-azido-TMMs (22), which were subsequently labelled with alkyne-containing biotin (DIBO-biotin) via click chemistry (23). Surface-exposed biotin-TMMs were recognized by Alexa Fluor 488-conjugated streptavidin, and visualized by fluorescence microscopy. (*B-E*) Representative bright-field and fluorescence microscopy images following DIBO-biotin/Alexa Fluor 488-streptavidin labelling of spheroplasts synthesizing (*B*) TMM, or 6-azido-TMM in the presence of (*C*) DMSO, (*D*) BM212 (2xMIC), and (*E*) AU1235 (2xMIC). Scale bars = 3 μm. (*F*) The fluorescence intensity per unit area for individual spheroplasts (N=100) in each condition (*B-E*) is plotted, with the medians and interquartile ranges indicated. Mann-Whitney test: ****, p < 0.0001 compared to the “no drug treatment” control.

To establish whether MmpL3 is responsible for flipping TMM across the IM, we employed specific MmpL3 variants that render *M. smegmatis* cells less sensitive to BM212, and tested if TMM flipping in spheroplasts expressing these variants would be more resistant to the effects of BM212. The growth of cells expressing MmpL3_V197M_ or MmpL3_A326T_ variants is only fully inhibited in the presence of 4-8 times the concentration of BM212 that inhibits wild-type growth (SI Appendix, Table S1) (9). We showed that BM212 does not affect MmpL3 levels in wild-type spheroplasts (SI Appendix, Fig. S7); yet it strongly inhibits TMM flipping in a dose-dependent manner (Fig. 5*A* and *D*). We further demonstrated that BM212 is less effective at reducing LysB accessibility to TMM in spheroplasts expressing MmpL3_V197M_ or MmpL3_A326T_ variants (Fig. 5*B*, *C* and *D*). In fact, BM212 is also unable to inhibit the display of 6-azido-TMMs on the surface of spheroplasts expressing MmpL3_V197M_ (Fig. 5*E*, *F*, *G* and *H*). Since TMM is only accessible on the outer leaflet of the IM in the presence of functional MmpL3 (i.e. not inhibited by BM212), we conclude that MmpL3 is the TMM flippase.

**Fig. 5.**
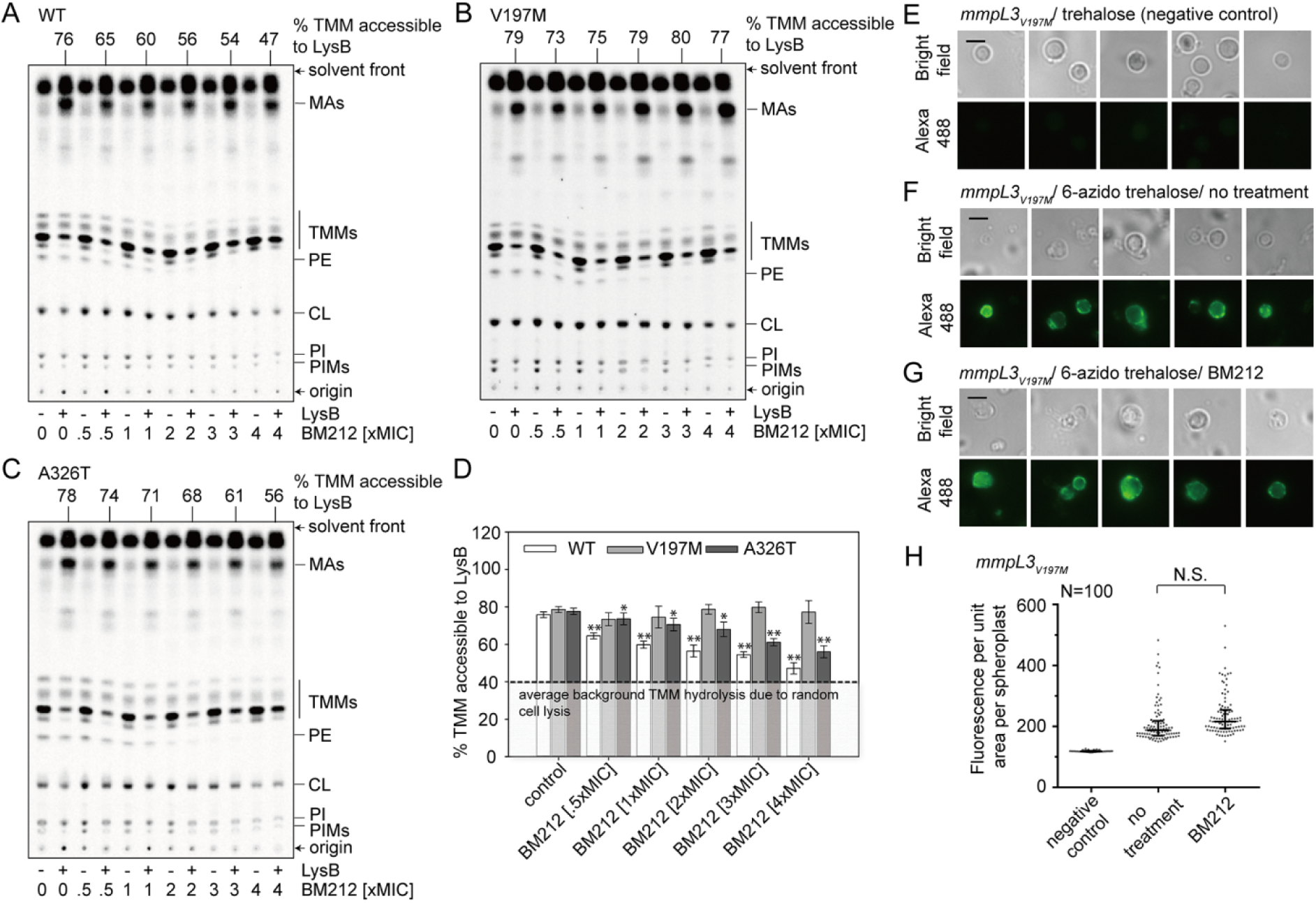
Mutations in MmpL3 render BM212 less effective in the inhibition of TMM flipping across the IM. (*A-C*) Representative TLC analyses of [^14^C]-labelled lipids newly-synthesized in the presence of indicated concentrations of BM212, and extracted from (*A*) WT, (*B*) *mmpL3*_*V197M*_, and (*C*) *mmpL3*_*A326T*_ *M. smegmatis* spheroplasts following treatment with or without purified LysB. Equal amounts of radioactivity were spotted for each sample. The developing solvent system comprises chloroform-methanol-water (30:8:1). (*D*) A graphical plot showing the dose-dependent effects of BM212 on the percentage of TMMs accessible to LysB in the respective spheroplasts (quantification as per Fig. 3). Average percentages and standard deviations from three biological replicates are plotted. The average background of TMM hydrolysis due to random cell lysis during the experiment (˜40%) is indicated. Student’s t-test: *, p < 0.05; **, p < 0.01 compared to the corresponding DMSO controls for each respective strain. (*E-G*) Representative bright-field and fluorescence microscopy images following DIBO-biotin/Alexa Fluor 488-streptavidin labelling of *mmpL3*_*V197M*_ spheroplasts synthesizing (*E*) TMM, or 6-azido-TMM in the presence of (*F*) DMSO, and (*G*) BM212 (2xMIC). Scale bars = 3 μm. (*H*) The fluorescence intensity per unit area for individual spheroplasts (N=100) in each condition (*E-G*) is plotted, with the medians and interquartile ranges indicated. Mann-Whitney test: N.S., p > 0.5 compared to the “no treatment” control.

### BM212 binds and directly inhibits MmpL3 function

MmpL3 function is believed to require the pmf, specifically the proton gradient (7). Consistently, we showed that proton gradient uncouplers, such as CCCP and nigericin (Fig. 3), but not membrane potential disruptors, such as valinomycin-K^+^ (SI Appendix, Fig. S8), can inhibit LysB accessibility to TMMs in spheroplasts. Whether BM212 inhibits TMM flipping by directly targeting MmpL3 is not clear. Contrary to previous reports (24), we did not observe effects on the proton gradient (SI Appendix, Table S2) nor the membrane potential (SI Appendix, Fig. S10) in spheroplasts treated with BM212, at concentrations that inhibited TMM flipping. Therefore, BM212 inhibits MmpL3 not via indirect effects on the pmf. To determine if BM212 directly targets MmpL3, we examined the ability of BM212 to bind physically to purified MmpL3 (SI Appendix, Fig. S11) in vitro. We demonstrated that [^14^C]-BM212 binds purified wild-type MmpL3 in a saturable manner (Fig. 6*A*). Furthermore, we showed that this interaction can be competed away by excess non-labelled BM212 (Fig. 6*B*), indicating that BM212 can bind specifically to MmpL3. In fact, mutations in *mmpL3* that confer resistance to BM212 mostly cluster in a specific region when mapped onto a structural model of MmpL3 (Fig. 6*C*), revealing a possible binding site. We conclude that BM212 inhibits TMM flipping across the IM by binding MmpL3 directly.

**Fig. 6.**
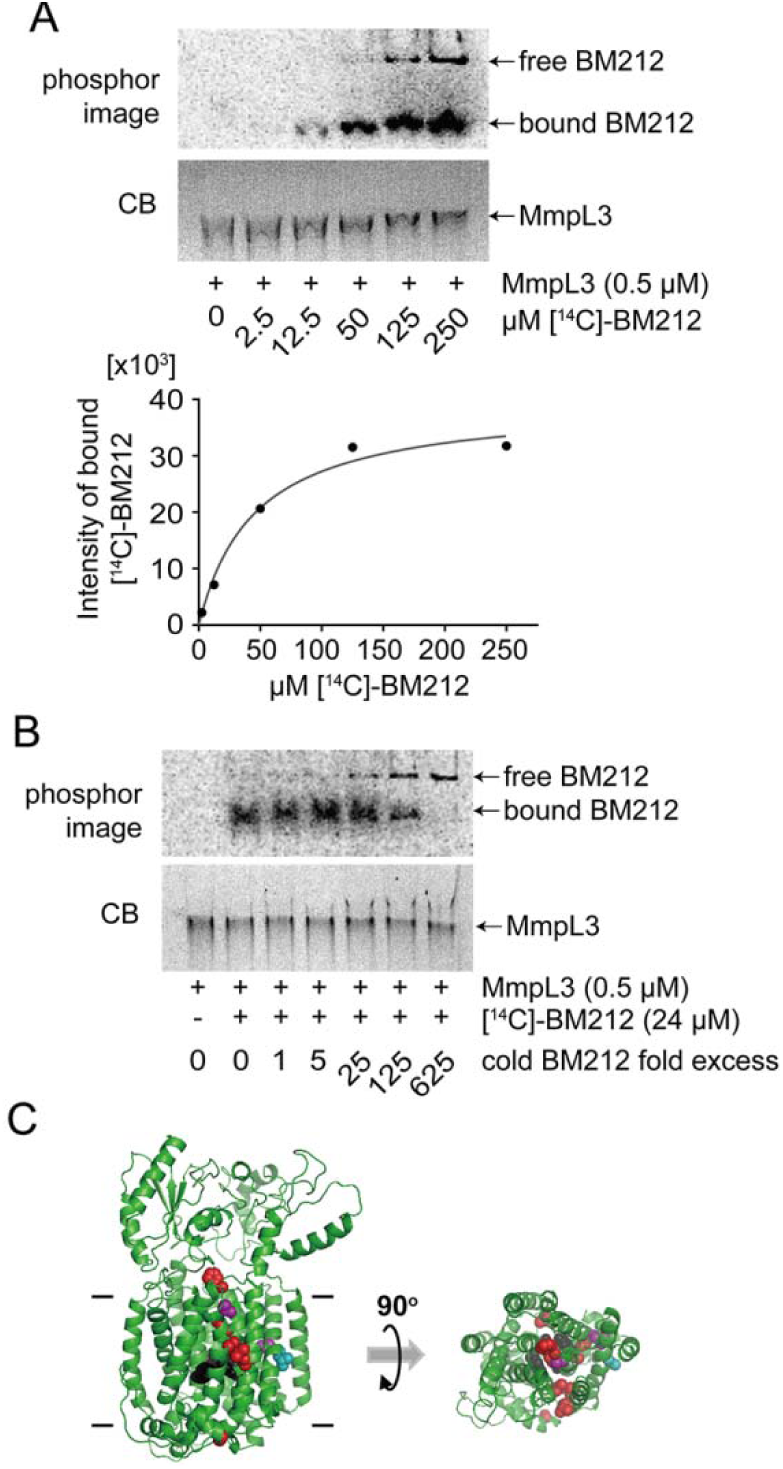
BM212 binds MmpL3 in vitro in a specific manner. (*A*) Clear Native-PAGE analyses of purified MmpL3-His samples incubated with increasing concentrations of [^14^C]-BM212, visualized separately by phosphor imaging and Coomassie blue (CB) staining. The amount of [^14^C]-BM212 bound to MmpL3 at each concentration was quantified and plotted. (*B*) Clear Native-PAGE analyses of purified MmpL3-His samples incubated with a fixed concentration of [^14^C]-BM212, but in the presence of increasing concentrations of cold BM212. Gels were visualized separately by phosphor imaging and CB staining. (*C*) Mutations that confer resistance against BM212 cluster on a structural model of MmpL3, suggesting a possible binding site. A Phyre2 (35) structural model for *M. smegmatis* MmpL3 without its C-terminal cytoplasmic domain is shown in side (*left*) and top (*right*) views. For clarity, periplasmic domains are removed from the top view image. Residues important for passage of protons are highlighted in black. Residues that conferred resistance against BM212 (10) when mutated in MmpL3 from *M. smegmatis*, *M. bovis BCG* and *M. tuberculosis* are highlighted in red, purple and cyan, respectively.

## Discussion

How the mycobacterial OM is assembled is not well understood. Many members of the MmpL protein family are believed to be transporters that contribute to the assembly of various OM lipids (25-28); however, their specific roles have not been clearly defined. MmpL3 is the only member of this family essential for growth (5-7). Using two independent assays that allow determination of TMM topology in the IM of mycobacterial spheroplasts, and employing putative inhibitors as molecular probes to modulate protein function, we have provided strong biochemical evidence that establish MmpL3 as the TMM flippase. Whether MmpL3 is the only protein mediating TMM flipping, or whether it is also involved in TMM release from the IM is not clear (Fig. 1*A*). One can posit that a second yet identified protein may be necessary for extracting TMM from the IM and handing it to a putative chaperone. This scenario would be comparable to the transport of lipoproteins across the cell envelope in Gram-negative bacteria (29). Alternatively, this step may also be mediated by MmpL3, in which case, flipping and release of TMM might be coupled; this would suggest MmpL3 could interact with the putative chaperone. Extending from this idea, a third scenario may be possible where TMM flipping and release are essentially one single step in a mechanism similar to RND efflux pumps, whereby TMM never really resides in the outer leaflet of the IM. This latter model is, however, less like since we have been able to decouple these steps by observing TMM translocation across the IM in our spheroplasts, which are effectively devoid of any putative chaperones. Moreover, despite being in the same RND protein superfamily, the structure of MmpL3 differs substantially from canonical RND efflux pumps (30). MmpL3 does form trimers like RND pumps (31), but the periplasmic domains of MmpL3 are much smaller (30), and it contains a large C-terminal cytoplasmic domain. Therefore, MmpL3 may not export TMM via an efflux mechanism. Further characterization of this system would be necessary to tease apart these models.

TB is one of the leading causes of death by infectious disease, and remains a major health problem worldwide (32). With the rapid emergence of multi-and extensive-drug resistant (MDR/XDR) TB, there is an urgent need to develop anti-TB drugs with novel mechanisms-of-action. In this regard, drugs inhibiting MmpL3, which has been shown to be an ideal target (33), would be especially important. While many small molecules are thought to inhibit MmpL3, it is puzzling how molecules with different molecular scaffolds may bind and target the same protein. We have now developed assays that measure the topology of TMM in the IM of mycobacterial spheroplasts, allowing the validation of true MmpL3 inhibitors. As a start, we have established that BM212 binds MmpL3 directly and inhibits its function. Furthermore, we have shown that SQ109, a molecule that has reached phase IIb clinical trials (Sequella), does not actually inhibit TMM flipping. In fact, it is likely that many of these molecules do not inhibit MmpL3, and have other targets, as has been shown for tetrahydropyrazo[1,5-*a*]pyrimidine-3-carboxamides (34). Our assays will help to select and advance small molecules currently under development as MmpL3-targeting drugs.

## Materials and Methods

Detailed descriptions of all experimental procedures can be found in the *SI Appendix*.

### Assessing TMM accessibility to degradation by purified LysB in spheroplasts. *M.*

*smegmatis* spheroplasts were metabolically labelled with sodium [1-^14^C]-acetate for 2 h, followed by addition of purified LysB for 30 min at 37°C. Lipids were extracted directly after the LysB treatment, analyzed by TLC, and visualized via phosphor imaging. Where indicated, putative MmpL3 inhibitors were added 15 min prior to addition of [^14^C]-acetate.

#### 6-azido-TMM surface display assay

*M. smegmatis* spheroplasts were metabolically labelled with 6-azido-trehalose for 2 h at 37°C to synthesize 6-azido-TMMs, which were then reacted with Click-IT^®^ Biotin DIBO alkyne to generate biotin-TMMs. Surface-exposed biotin-TMMs were detected on spheroplasts using Alexa Fluor ^®^ 488-conjugated streptavidin, and visualized by fluorescence microscopy. Where indicated, putative MmpL3 inhibitors were added 15 min prior to addition of 6-azido-trehalose, and included in all wash buffers.

#### BM212-MmpL3 binding assay

Purified MmpL3 was incubated with indicated concentrations of [^14^C]-BM212 and/or cold BM212 for 30 min at room temperature. Samples are analyzed using Clear Native-PAGE, followed by phosphor imaging and CB staining.

## Acknowledgments

We thank Graham Hatfull (U Pittsburgh) for providing the pLAM3 plasmid for LysB overexpression, and Derek Lin (NUS) for constructing the inactive LysB_S82A_ variant. We are grateful to Benjamin Swarts (Central Michigan U) for his generous gift of 6-azido-trehalose. We also thank Eric Rubin (Harvard School of Public Health) for providing strains, and for critical discussion and comments on the manuscript. We acknowledge Jasmine Chen (Mechanobiology Institute) for help with microscopy. This work was supported by the National University of Singapore Start-up funding, the Singapore Ministry of Education Academic Research Fund Tier 1 and Tier 2 (MOE2014-T2-1-042) grants (to S.-S.C.).

## Classification

Biological Sciences - Biochemistry

